# Atomic resolution cryo-EM at 200 keV

**DOI:** 10.64898/2026.01.09.698548

**Authors:** Radostin Danev, Haruaki Yanagisawa, Keitaro Yamashita, Fabian Eisenstein, Masahide Kikkawa

## Abstract

Atomic resolution in cryo-electron microscopy was first demonstrated six years ago. This was accomplished using 300 kV electron microscopes equipped with new hardware that provided narrower energy spread, aberration correction, and energy filtering. Here, we report the achievement of 1.24 Å atomic resolution on an upgraded 200 kV electron microscope featuring a cold field emission gun, a high-resolution objective lens polepiece, and an energy filter. These components transform the instrument into a cost-effective single particle cryo-EM platform with performance comparable to that of significantly more expensive 300 kV systems. The microscope can also be operated at 100 kV and by using a high-speed hybrid-pixel detector we were able to reach sub-2 Å resolution.

## 1. Introduction

The cryo-EM “resolution revolution” has been ongoing for more than a decade. It reached a major milestone in 2020 with the demonstration of atomic resolution by two independent teams (Nakane *et al*., 2020; Yip *et al*., 2020). According to the current consensus in X-ray crystallography, “atomic resolution” is defined as 1.2 Å or better (Dauter, 2003), based on a criterion of non-overlapping atomic densities proposed by Sheldrick (Sheldrick, 1990).

The first atomic resolution cryo-EM results were obtained using well-behaved, highly symmetric heavy-chain apoferritin, which has served as the gold standard test specimen for cryo-EM single-particle performance since its introduction by Russo and Passmore (Russo & Passmore, 2014) and its subsequent improvement to a heavy-chain variant (Danev *et al*., 2019). The two atomic resolution teams used upgraded 300 kV microscopes: one featured a cold field-emission gun (CFEG) and a newly developed energy filter (Nakane *et al*., 2020), and the other a monochromator and a spherical aberration corrector (Yip *et al*., 2020). Since these reports, other groups have also presented atomic resolution results on 300 kV instruments, almost all equipped with CFEG sources (Fujita *et al*., 2023; Zhang *et al*., 2020; Maki-Yonekura *et al*., 2023; Küçükoğlu *et al*., 2024). In 2020, we tested the newly installed Titan Krios G4 (Thermo Fisher Scientific, Waltham, USA) 300 kV electron microscope at the University of Tokyo equipped with a Schottky FEG (SFEG) and the dataset reached 1.31 Å (Danev *et al*., 2021). At present, 300 kV microscopes offer the best cryo-EM performance for both single-particle analysis and cryo-tomography. However, they are expensive to purchase, costly to maintain, heavy in electricity consumption, and require a large installation space.

The cryo-EM capabilities of 200 kV microscopes have been extensively evaluated over the years with remarkable results (Herzik *et al*., 2017; Wu *et al*., 2020; Merk *et al*., 2020; Hamdi *et al*., 2020; Kayama *et al*., 2021; Feathers *et al*., 2021; Gerle *et al*., 2022; Thangaratnarajah *et al*., 2022; Koh *et al*., 2022; Jia *et al*., 2024). Surprisingly, despite their excellent performance, only ∼10 % of deposited maps with resolutions better than 6 Å in the Electron Microscopy Data bank (EMDB) come from 200 kV microscopes. Currently, 200 kV machines are widely used as a cost-effective alternative to 300 kV instruments at universities, research institutions, and industrial facilities for sample screening and data collection, with the understanding that the achievable resolution may not be as high but is often sufficient to answer most research questions.

In the past few years, 100 kV microscopes were shown to hold substantial promise as “people’s cryo-microscopes”, offering sufficient performance for sample screening and exploratory data collection at an affordable cost and with minimal installation requirements (Naydenova *et al*., 2019; McMullan *et al*., 2023; Chan *et al*., 2024; Venugopal *et al*., 2025; Karia *et al*., 2025). However, achieving high resolution results on current commercially available 100 kV instruments remains challenging, as evidenced by the fact that there are fewer than fifty maps at resolutions better than 6 Å in the EMDB, the majority of which are test samples.

Here, we evaluated the performance of an upgraded 200 kV microscope equipped with a CFEG, a narrow gap objective lens polepiece, and an omega-type energy filter. We collected apoferritin test datasets at accelerating voltages of 200 and 100 kV using a latest generation direct detection camera and a high-speed hybrid-pixel detector respectively. The results are presented below.

## 2. Materials and methods

### 2.1 Microscope configuration

The experiments were performed on a CRYO ARM 200 II (JEM-Z200CA, JEOL Ltd., Tokyo, Japan) electron cryo-microscope upgraded with a narrow-gap high-resolution objective lens polepiece. This reduced the spherical and chromatic aberration coefficients from the regular *C_S_* = 2.7 mm and *C_C_* = 2.8 mm to *C_S_* = 1.5 mm and *C_C_* = 1.8 mm. The microscope is equipped as standard with a CFEG, a three-lens condenser system, and an omega-type energy filter. The system also featured a Gatan K3 (Gatan, Pleasanton, USA) direct electron detector and an electrostatic dose modulator (EDM), which was used as a fast electrostatic shutter. For the 100 kV experiments, the camera was replaced with a DECTRIS SINGLA (DECTRIS Ltd., Baden-Daettwil, Switzerland) high-speed hybrid-pixel detector.

### 2.2 Sample preparation

Mouse heavy chain apoferritin was expressed and purified as described previously (Danev *et al*., 2021). Cryo-EM samples were prepared by applying 3 μl of 1.9 mg/ml sample solution on UltrAuFoil R0.6/1 300 mesh (200 kV experiments) or UltrAuFoil R1.2/1.3 300 mesh (100 kV experiments) grids (Russo & Passmore, 2016) (Quantifoil Micro Tools GmbH, Jena, Germany) and plunge-freezing in liquid ethane on a Vitrobot Mark IV (Thermo Fisher Scientific, Waltham, USA), blot time 20 s (R0.6/1.0 grids) or 10 s (R1.2/1.3 grids), chamber temperature 4 °C, 100 % humidity.

### 2.3 Data collection

The datasets were collected automatically by SerialEM software (Schorb *et al*., 2019) using its built-in single-particle automation routines. The main acquisition parameters are summarized in Supplementary Table S1.

For the 200 kV dataset, the microscope was set up at an indicated magnification of 150,000x, calibrated pixel size 0.3056 Å pixel^-1^, spot size 3, convergence angle 3, condenser aperture 100 μm, beam diameter ∼0.95 μm, no objective aperture, zero-loss energy filtering with 20 eV slit, target defocus -0.5 μm. The detector was operated in counting (non-super-resolution) correlated double sampling (CDS) mode, with an exposure time of 1.51 s, an exposure rate of 3.3 e pixel^-1^ s^-1^, a total exposure of 53.4 e Å^-2^, a frame time of 0.0185 s, 81 frames, and an exposure per frame of 0.66 e Å^-2^. In total, 13,654 movies were collected in 56 hours with an average throughput of 244 movies per hour using a beam-tilt compensated 3x3x1 image shift acquisition pattern with a single image in the center of each hole.

For the 100 kV dataset, the microscope was set up at an indicated magnification of 500,000x, calibrated pixel size 1.17 Å pixel^-1^, spot size 6, convergence angle 3, condenser aperture 70 μm, beam diameter ∼0.53 μm, objective aperture 250 μm, zero-loss energy filtering with 20 eV slit, and target defocus -0.45 μm to -0.55 μm. The movies were recorded in HDF5 format at the raw framerate of the detector of 4,500 frames s^-1^, with an exposure time of 3.0 s, an exposure rate of 23.8 e pixel^-1^ s^-1^, a total exposure of 52.2 e Å^-2^, 13,500 total frames, an exposure per frame of 0.0039 e Å^-2^. In total, 17,424 movies were collected in 49.5 hours with an average throughput of 352 movies per hour using a beam-tilt compensated 3x3x4 image shift acquisition pattern with four images in each hole.

### 2.4 Data processing

The data were processed with CryoSPARC (Punjani *et al*., 2017) (Structura Biotechnology, Toronto, Canada).

The 200 kV dataset processing workflow is summarized in Supplementary Fig. S1. Briefly, 13,654 movies were subjected to patch motion correction and patch CTF estimation followed by exposure curation with CTF fit resolution selection range between 2 and 4 Å, retaining 10,189 micrographs. The micrographs were split into nine exposure groups based on their image shift. Particles were picked with 20 Å low-pass filtered templates generated from a previous 3D apoferritin map, resulting in 927k picks after selection with NCC score > 0.4 and local power between 600 and 900. Extraction with a 160 pixel box at 1.57 Å pixel^-1^ produced 652k particles that were subjected to 2D classification with 100 classes. Manual 2D class selection left 635k particles that were run through ab-initio 3D reconstruction with two classes. The 3D reference and 622k particles from the larger class were used for further processing. Initial 3D homo refinement hit Nyquist at 3.23 Å. The particles were re-extracted with 400 pixel box at 0.627 Å pixel^-1^ and another 3D homo refinement reached 1.56 Å. The particles were then split into hour-wise exposure groups for a total of 504 exposure groups, followed by a 3D homo refinement which reached 1.35 Å. A round of reference-based motion correction and 3D homo refinement improved the map to 1.26 Å. A 3D heterogeneous refinement with three classes reduced the particle stack to 615k, followed by another round of reference-based motion correction and 3D homo refinement to produce the final map at 1.24 Å.

The 100 kV dataset processing workflow is summarized in Supplementary Fig. S2. Briefly, the 17,654 HDF5 hardware frame stacks were subjected to 2x super-resolution electron counting (Zambon, 2023) with a GPU-accelerated software tool provided by DECTRIS (DECTRIS Ltd., Baden-Daettwil, Switzerland) and were fractionated with 225 hardware frames per fraction into 60 frame MRC movies. The movies were renamed with a Python script according to image shift group specifications in summed MRC images that were saved separately by SerialEM during acquisition. The renamed super-resolution MRC movies were imported into CryoSPARC and were subjected to patch motion correction with 3/4 Fourier crop and patch CTF estimation followed by exposure curation with CTF fit resolution selection range between 2.5 and 5 Å, retaining 9,123 micrographs. Particles were picked with 20 Å low-pass filtered templates generated from a previous 3D apoferritin map, resulting in 477k picks after selection with NCC score > 0.4 and local power between 8*10^6^ and 13*10^6^. Extraction with a 150 pixel box at 1.4 Å pixel^-1^ produced 350k particles that were subjected to 2D classification with 200 classes. Manual 2D class selection left 322k particles that were run through ab-initio 3D reconstruction with three classes. The 3D reference and 321k particles from the largest class were used for further processing. Initial 3D homo refinement reached 2.9 Å. The particles were re-extracted with a 250 pixel box at 0.936 Å pixel^-1^ and split into 36 exposure groups according to image shift. Another 3D homo refinement reached 2.01 Å. A reference-based motion correction was attempted but did not improve the resolution while exhibiting unexpectedly smooth particle tracks and an anomalous overweighting of high-resolution spectral components in later movie frames. To circumvent that, patch motion correction of the movies was performed again by using only the first 30 frames. Particles were re-extracted from the limited-frame micrographs and a 3D homo refinement reached 2.0 Å. A round of reference-based motion correction and 3D homo refinement improved the map resolution to 1.93 Å. Splitting the particles into hour-wise exposure groups and further exposure curation reduced the particle stack to 250k particles and improved the 3D homogeneous refinement resolution to 1.91 Å.

To generate the Rosenthal-Henderson plot (Rosenthal & Henderson, 2003) for the 200 kV dataset, random particle subsets from the final particle stack were 3D homo refined independently and the results were plotted in a reciprocal squared resolution versus logarithm of the number of particles. The B-factor was calculated as 2*(slope of linear fit)^-1^ from the linear regression fits through the data points in these coordinates.

Evaluation of the effect of radiation damage by pre-exposure on the achievable resolution was performed by using the first 1,000 movies containing 58k particles from the 200 kV dataset, running motion correction jobs with omission of a varying number of initial movie frames (Supplementary Table S2), extracting the particles from the aligned micrographs, and performing a 3D homogeneous refinement with defocus, nine image-shift exposure group beam tilt, and Ewald sphere correction. Reference-based motion correction and higher-order aberration refinements were not used during this processing.

### 2.5 Numerical estimation of the effect of pre-exposure on resolution

To quantify the resolution decrease as a function of pre-exposure, we must estimate the spectral signal-to-noise ratio (SNR) and dampen it with a resolution-dependent radiation damage model. It is not necessary to know the absolute but rather the relative spectral SNR, which can be calculated from the B-factor fit of the Rosenthal-Henderson plot. Assuming that particles represent independent measurements, the SNR is inversely proportional to the square root of the number of particles necessary to reach a given resolution. Therefore, having a 3D refinement without pre-exposure, the relative SNR of lower resolution shells can be calculated as the square root of the particle number ratio. Then, by applying a resolution-dependent radiation damage model, the exposure needed to attenuate the relative SNR of a given resolution shell to 1 can be estimated. This will correspond to the exposure that will limit the 3D reconstruction to the resolution of that shell.

Using this approach and the empirical radiation damage model by Grant and Grigorieff (Grant & Grigorieff, 2015) adjusted for 200 kV by reducing the critical exposure by 25% (as suggested in the paper), we estimated the resolution vs pre-exposure. The calculations were performed in Wolfram Mathematica (Wolfram Research, Champaign, USA).

### 2.6 Model refinement and visualization

The initial model (PDB-7A4M) was rigid-body fitted in Coot (Casañal *et al*., 2020), followed by reciprocal-space refinement using Servalcat (Yamashita *et al*., 2021) against unsharpened half maps. The model was interactively adjusted in Coot using the sharpened map and *F_o_ − F_c_* difference map, which included the positioning of polar hydrogen atoms. Finally, a hydrogen omit *F_o_ − F_c_* difference map was calculated using Servalcat. The models were validated using MolProbity (Chen *et al*., 2010).

3D images of the maps and model for the figures were created using UCSF Chimera (Pettersen *et al*., 2004) and UCSF Chimera X (Meng *et al*., 2023).

## 3. Results

Fig. 1 shows normalized histograms of the resolution of Electron Microscopy Data Bank (EMDB) depositions from 300 kV and 200 kV microscopes in the last six years. The ratio of 200 kV to 300 kV depositions is approximately 1:9 (3,058 vs 27,825). Log-normal fits of the histograms indicate a 0.25 Å higher median resolution at 300 kV. The rather moderate resolution advantage on its own cannot explain the big disparity in the number of depositions. Nevertheless, a closer look at the histograms shows that 300 kV is ∼1.7 times more prolific in the 2.5–3.0 Å range, and ∼3.3 times more prolific in the 2.0–2.5 Å range. These resolution ranges, and especially the latter, are of increasing importance in cryo-EM studies because they enable more accurate modeling of sidechains, identification and pose assignment of small molecules, and high-confidence detection of water molecules.

**Figure 1.**
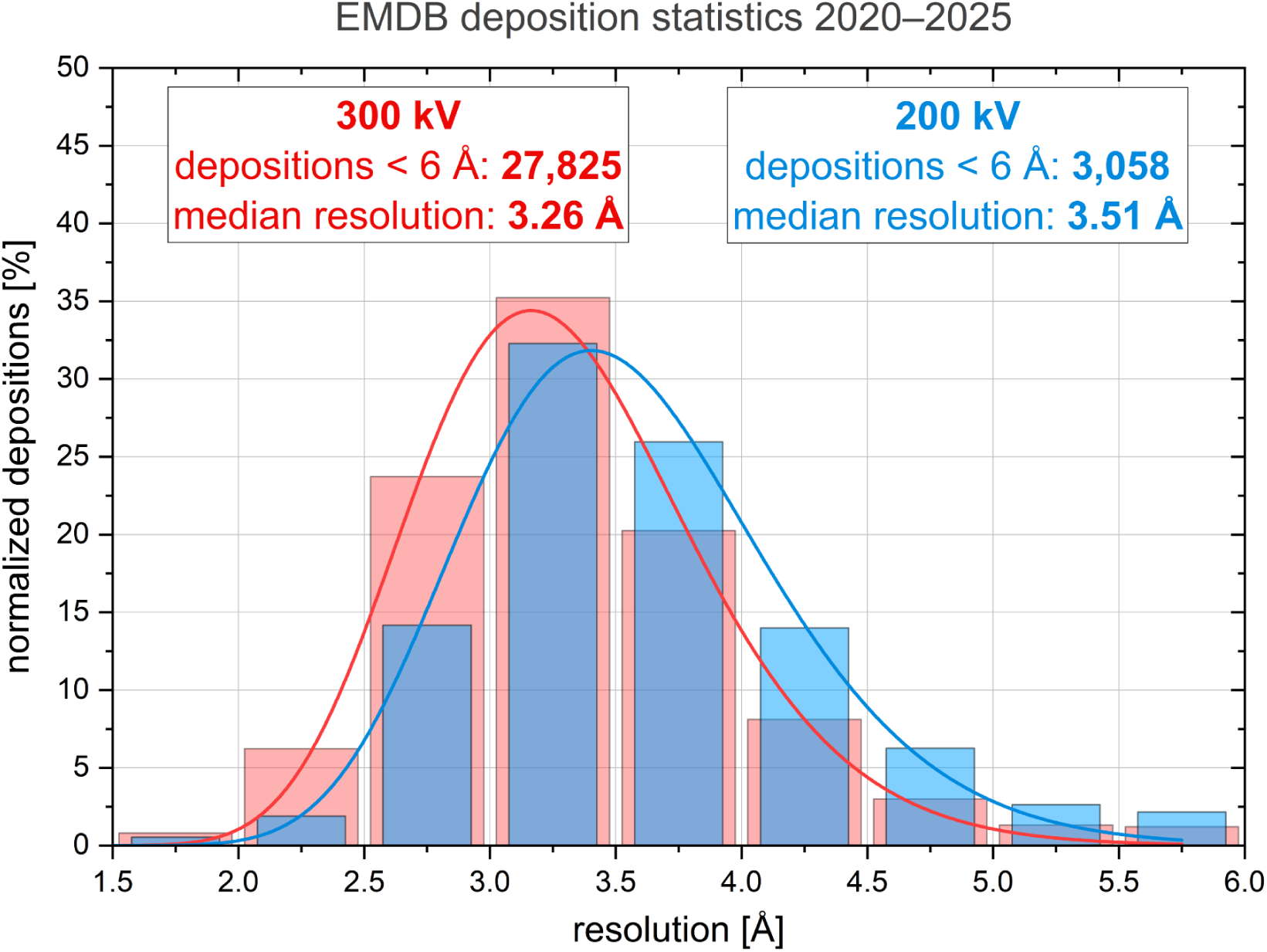
Resolution statistics of cryo-EM maps with resolution below 6 Å from 300 kV and 200 kV microscopes deposited in the Electron Microscopy Data Bank (EMDB) in the last six years (2020–2025). The resolution histograms (pink and light blue bars) were normalized by the number of depositions. Log-normal fits of the histograms are shown with red and blue lines. The number of depositions and median resolution from the log-normal fits are listed in the text boxes.

Fig. 2 contains a plot of the theoretical temporal coherence contrast transfer function (CTF) envelope (eq. 6.42 in (Reimer & Kohl, 2010)) for a number of cryo-microscope configurations. The top-of-the-line CRYO ARM 300 II (JEOL Ltd., Tokyo, Japan) and Krios 5 (Thermo Fisher Scientific, Waltham, USA) (green solid line) 300 kV machines have demonstrated atomic resolution in the past (Nakane *et al*., 2020; Yip *et al*., 2020; Maki-Yonekura *et al*., 2023) and are currently the highest optical performance commercially available cryo-microscopes. The CRYO ARM 200 II (pink solid line) used in this study is a close second and is currently the top performing 200 kV instrument. Other CFEG 200 kV microscopes, such as the CRYO ARM 200 (JEOL Ltd., Tokyo, Japan) and Glacios 2 (Thermo Fisher Scientific, Waltham, USA) (blue solid line), follow next. Remarkably, the recently presented “Dublin lens” design for 100 keV (Alves *et al*., 2025) (gray solid line) with a very low chromatic aberration coefficient of *C_C_*= 1.0 mm, if coupled with a CFEG will perform on par with the CRO ARM 200 and Glacios 2. SFEG-equipped instruments, such as the Krios, Glacios (Thermo Fisher Scientific, Waltham, USA), the “Dublin lens”, and Tundra (Thermo Fisher Scientific, Waltham, USA) come next in terms of theoretical optical quality.

**Figure 2.**
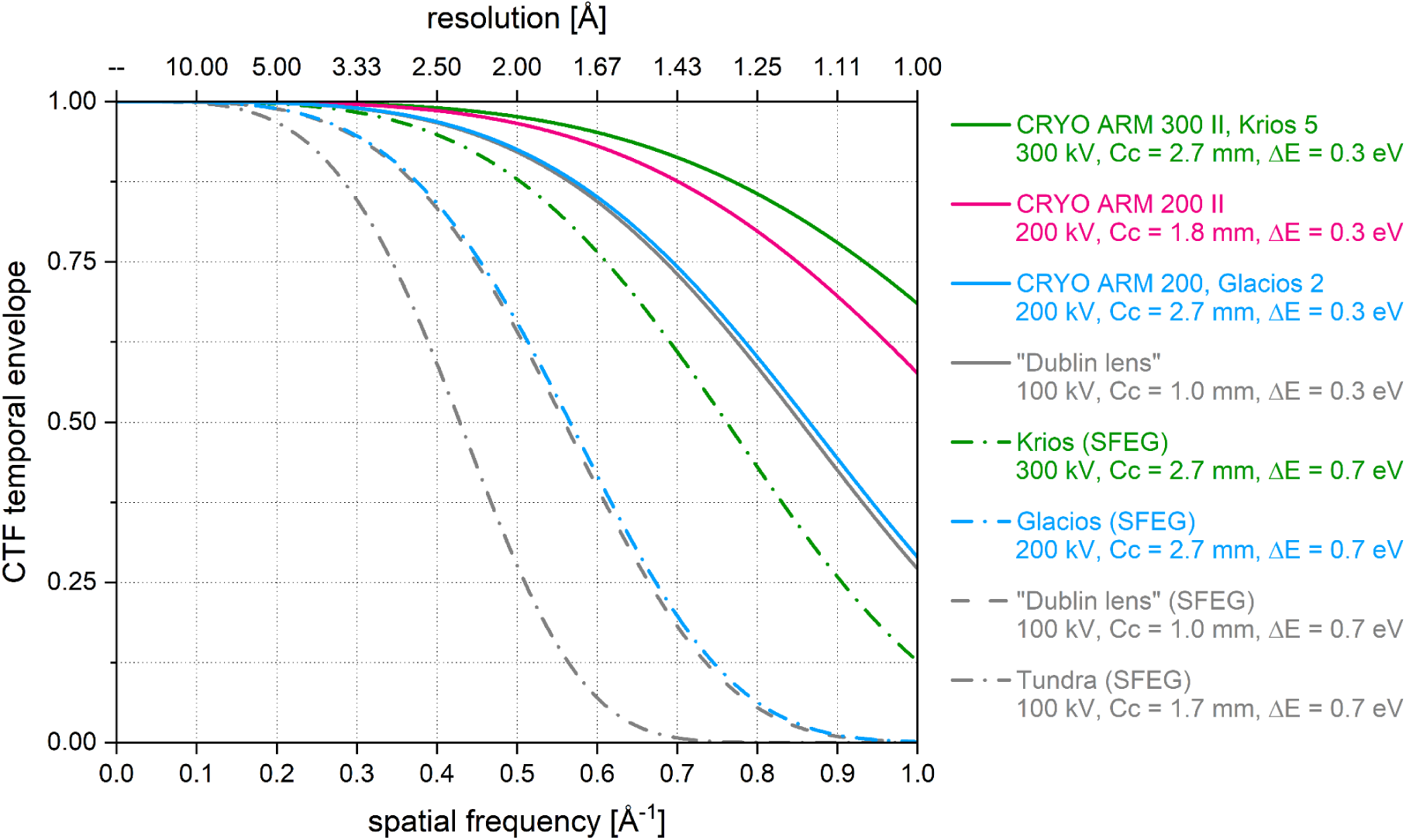
Theoretical temporal coherence envelope of various microscope configurations. Solid lines represent Cold Field Emission Gun (CFEG) instruments (ΔE = 0.3 eV). Dashed and dot-dashed lines correspond to Schottky (thermionic) FEG machines (ΔE = 0.7 eV). The CRYO ARM 200 II (pink line) evaluated here is second in optical performance after the top-of-the-line CRYO ARM 300 II and Krios 5.

To experimentally test the overall optical performance of the CRYO ARM 200 II, we collected high-magnification (200,000x) images of oriented single crystal (100) gold test specimen (AGS135, Agar Scientific, Rotherham, UK). The Fourier transform of one of the best images is shown in Fig. It contains spots up to lattice plane (046) with spacing 0.566 Å along the sample stage tilt axis and up to lattice plane (044) with spacing 0.721 Å perpendicular to the tilt axis. This confirms the excellent optical performance of the microscope going beyond 1 Å in all directions. It also indicates that it may have been slightly affected by mechanical disturbances. The instrument is installed in a room that is on the ground floor of a building but was not specially designed for high-resolution electron microscopy with considerations and measures to minimize floor vibrations.

The data acquisition scheme and exemplary micrograph from the 200 kV apoferritin dataset are presented in Figs. 4a, b. To minimize beam-induced sample motion we prepared the 200 kV sample on 0.6 μm holes gold foil grid and collected a single image in the center of each hole with the beam illuminating uniformly the hole edge all-around (Fig. 4a). To maximize the camera’s detective quantum efficiency (DQE) performance over the target resolution range, the 200 kV data was collected at a relatively high indicated magnification of 150,000x with a pixel size of 0.3056 Å pix^-1^ (Fig. 4b, Supplementary Table S1). The resolution of the final 3D map corresponds to ∼49% of the physical Nyquist limit of the camera.

**Figure 3.**
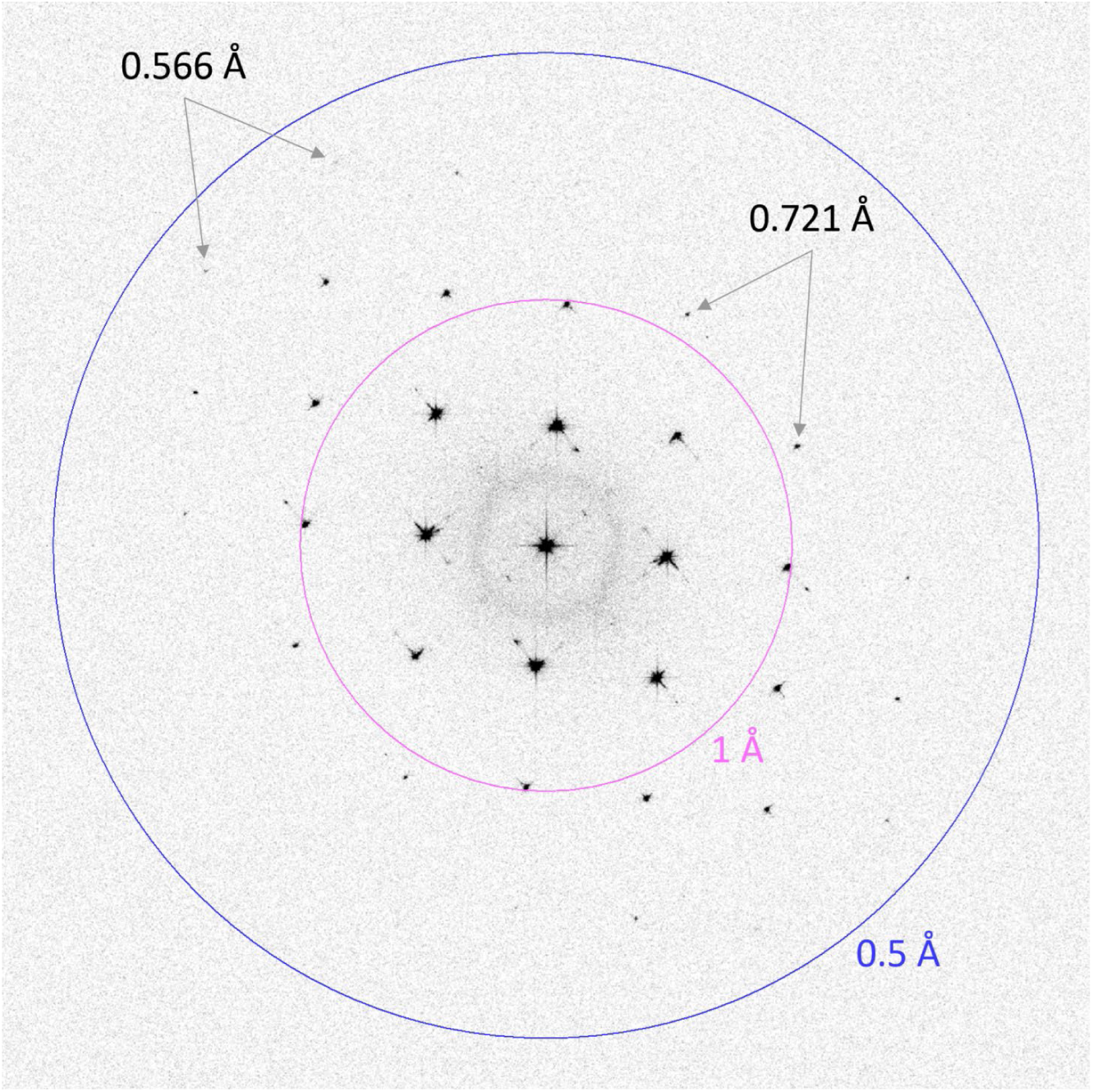
2D Fourier transform amplitudes of an oriented single crystal (100) gold sample image from the CRYO ARM 200 II. The visibility of lattice plane (046) spots corresponding to 0.566 Å spacing in the direction of the sample stage tilt axis and lattice plane (044) spots corresponding to 0.721 Å spacing in the perpendicular direction attest to the excellent optical performance of the microscope, going beyond 1 Å in all directions.

**Figure 4.**
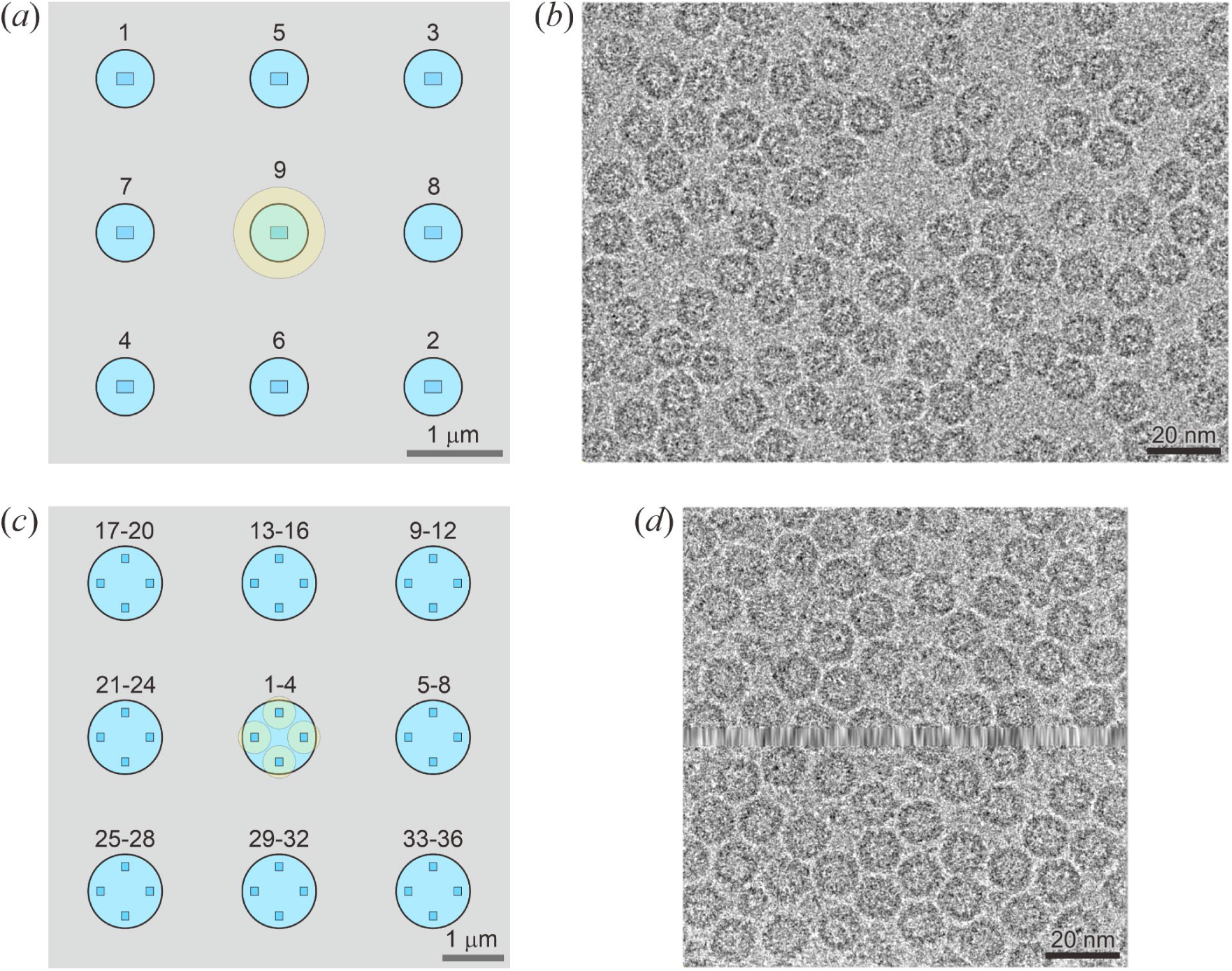
Acquisition patterns and exemplary images for the (*a, b*) 200 kV and (*c, d*) 100 kV apoferritin test datasets. (*a, c*) Acquisition pattern schematics with support film holes in light blue, detector size and shape in blue, and beam size and position in pale yellow. Numbers indicate the acquisition order. (*b, d*) Exemplary images from the datasets collected at (*b*) 200 kV on a Gatan K3 camera, and (*d*) 100 kV on a DECTRIS SINGLA hybrid-pixel detector. The interpolated inter-module gap of the SINGLA detector is visible as a blurred horizontal stripe through the middle of the image.

The 100 kV dataset was also collected on a gold foil grid but to maximize data acquisition throughput we used a grid with 1.2 μm holes and acquired four images per hole (Fig. 4c). In previous experiments using such acquisition strategy on this and other microscopes, we have reached resolutions far below 2 Å (Danev *et al*., 2021) and therefore do not expect it to be a limiting factor in this test. Due to the much larger physical pixel of the DECTRIS SINGLA camera (75 μm vs 5 μm of the Gatan K3) we had to collect at even higher magnification of 500,000x (Supplementary Table S1). This presented some practical challenges in terms of microscope alignment and operation in the ultra-high magnification range, which is not typically used in cryo-EM. Fig. 4d contains an exemplary image from the 100 kV dataset illustrating the gap (horizontal stripe in the middle of the image) between the two modules of the detector which is interpolated in software to facilitate cryo-EM data processing.

The results from the processing of the 200 kV dataset are presented in Fig.5. The reconstruction (Fig. 5a) reached 1.24 Å according to the 0.143 gold-standard Fourier Shell Correlation (FSC) criterion (Fig. 5b). The 3D map shows features consistent with the estimated resolution, such as atomic bulges at a lower surface threshold and individual blobs for non-hydrogen atoms at a higher threshold (Fig.5c, gray and blue surfaces). Furthermore, an *F_o_–F_c_* difference density map calculated in Servalcat (Yamashita *et al*., 2021) revealed hydrogen atom densities at many of their expected positions (Fig. 5c, green surfaces).

**Figure 5.**
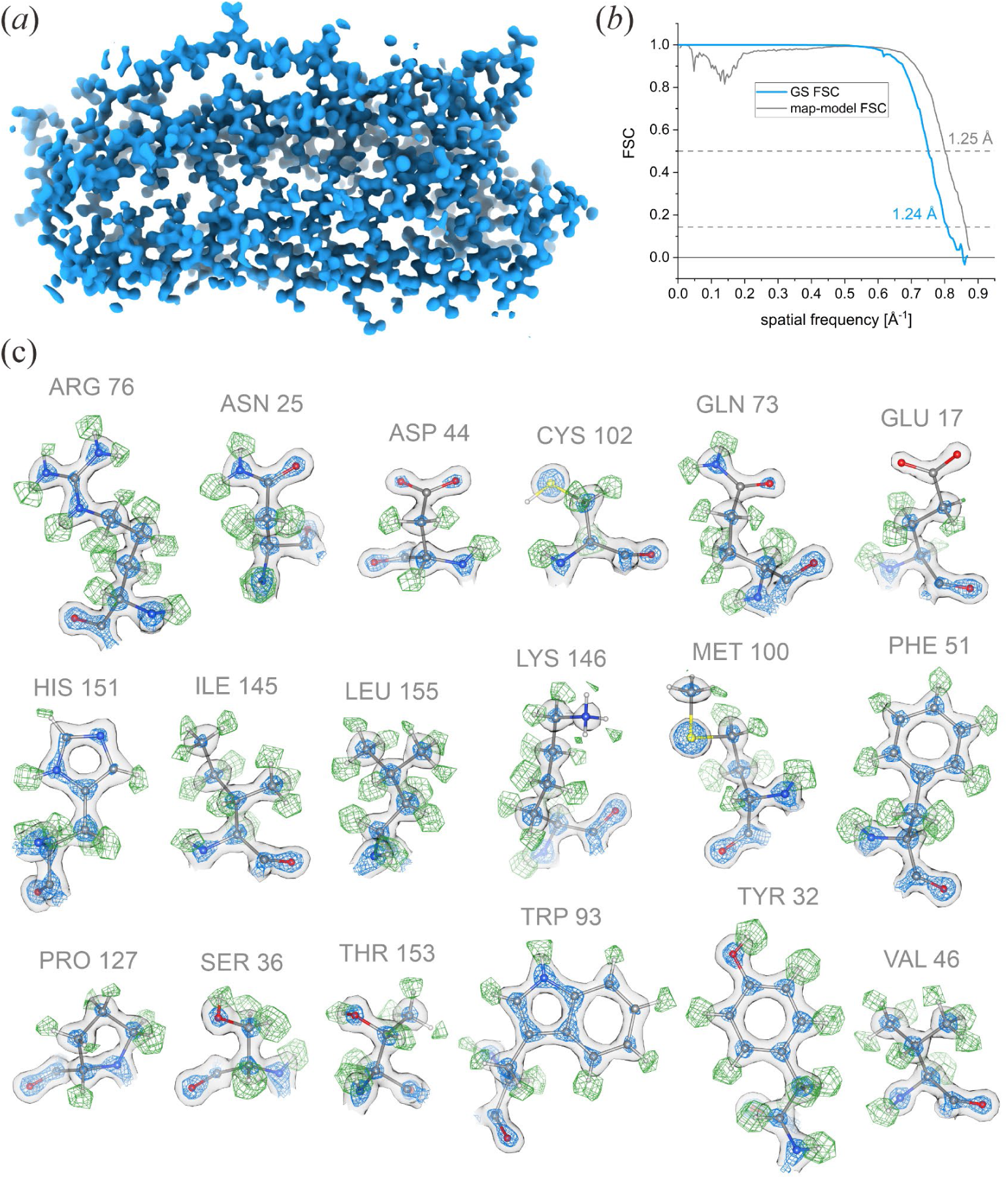
Atomic resolution cryo-EM reconstruction of the 200 kV apoferritin dataset. (*a*) Overview of the 3D density map of a monomer. (*b*) Plot of the gold-standard FSC (GS FSC) indicating 1.24 Å resolution at 0.143 level and the map-model FSC indicating 1.25 Å resolution at 0.5 level. (*c*) Sidechain map features. The gray surface is at a lower threshold level typically used to depict the map. The blue surface is at a high threshold level, showcasing the atomic resolution of the map with blobs for individual atoms. The green surface represents an *Fo - Fc* difference density map (3σ level, normalized within a mask) calculated from a hydrogen-omit model. Positive density peaks are clearly visible at positions corresponding to hydrogen atoms in the molecular model.

We quantified the overall experimental performance using a Rosenthal-Henderson B-factor plot (Rosenthal & Henderson, 2003). Fig. 6 contains B-factor plots for the 200 kV dataset (red symbols and line) and the Krios G4 (set 1 in (Danev *et al*., 2021)) apoferritin dataset (blue symbols and line). The measured B-factors were 39.0 Å^2^ for the CRYO ARM 200 II and 38.7 Å^2^ for the Krios G4. In practical terms, they are virtually identical. The CRYO ARM 200 II plot was offset vertically towards slightly higher resolution which indicates a slightly higher overall signal-to-noise ratio of the data.

**Figure 6.**
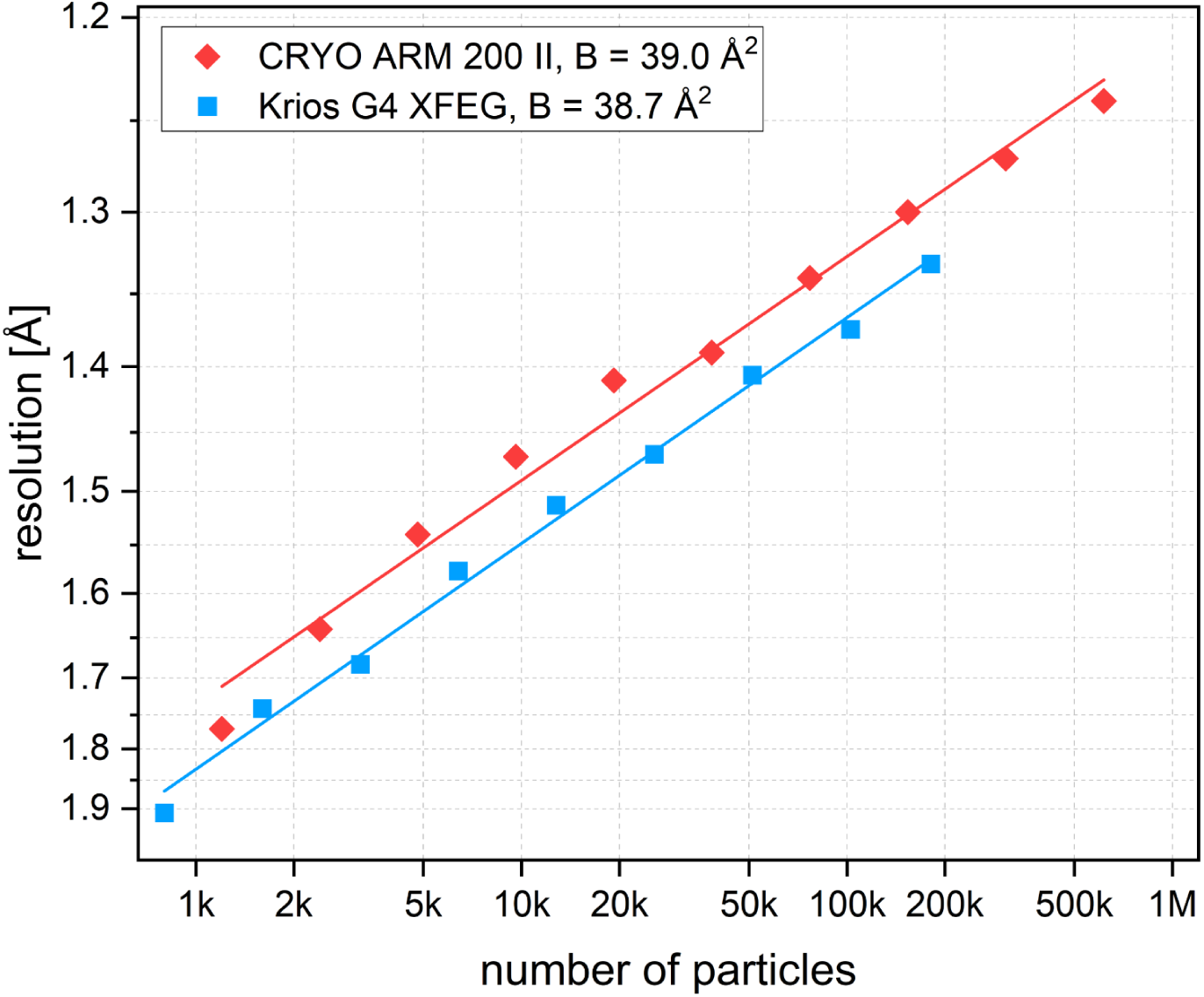
Rosenthal-Henderson B-factor plots comparing the overall cryo-EM performance of the CRYO ARM 200 II (red symbols and line) and the Krios G4 (blue symbols and line) microscopes with apoferritin samples. The slopes of the linear fits (red and blue lines) correspond to B-factors of 39.0 Å^2^ and 38.7 Å^2^, respectively.

During data processing, we observed a 0.2 Å improvement in resolution (from 1.56 Å to 1.35 Å) after splitting the data into hour-wise exposure groups (Supplementary Fig. S1), indicating that there was a variation over time of the optical parameters. Supplementary Fig. S3 shows a temporal trend plot of the refined beam tilt for the nine image-shift exposure groups. The beam tilt steadily increased by ∼0.65 mrad over the course of the experiment. It is not clear what caused this change. There were no significant variations in the room and cooling water temperatures. One possible explanation is a gradual drift of the optics because the lens degaussing routines were disabled to reduce time overhead. The plot also shows that SerialEM’s image shift beam tilt-compensation performed well by keeping the spread between the nine acquisition areas within 0.1 mrad.

We were curious about the effect of radiation damage on the achievable resolution. To quantify it, we used the first 1,000 movies and performed independent reconstructions starting from motion correction by omitting a varying number of initial frames to simulate pre-exposure. The same set of particle coordinates was used to extract particles from the aligned micrographs, followed by a 3D homogeneous refinement job. The results are presented in Fig. 7. Surprisingly, the reconstruction reached ∼2.9 Å resolution even after 20 e Å^-2^ of pre-exposure. The resolution dependence on pre-exposure was fitted very well (*R*^2^ = 0.997) by a parabola (Fig.7a, red line). Numerical estimation of the resolution vs pre-exposure based on applying an empirical radiation damage model (Grant & Grigorieff, 2015) produced results that underestimated (larger value) the achievable resolution (Fig. 7a, black symbols). Using a total exposure value that was 82 % of the experimentally measured one in the numerical calculation produced results that were much closer to the experimental ones (Fig. 7a, blue symbols). The difference in the exposure parameter could be due to various reasons, such as empirical damage model accuracy for this particular sample and support film, variation of the actual exposure during the experiment due to CFEG current decline, or other factors. Fig. 7b contains panels illustrating the decline of 3D map fidelity with increasing pre-exposure.

**Figure 7.**
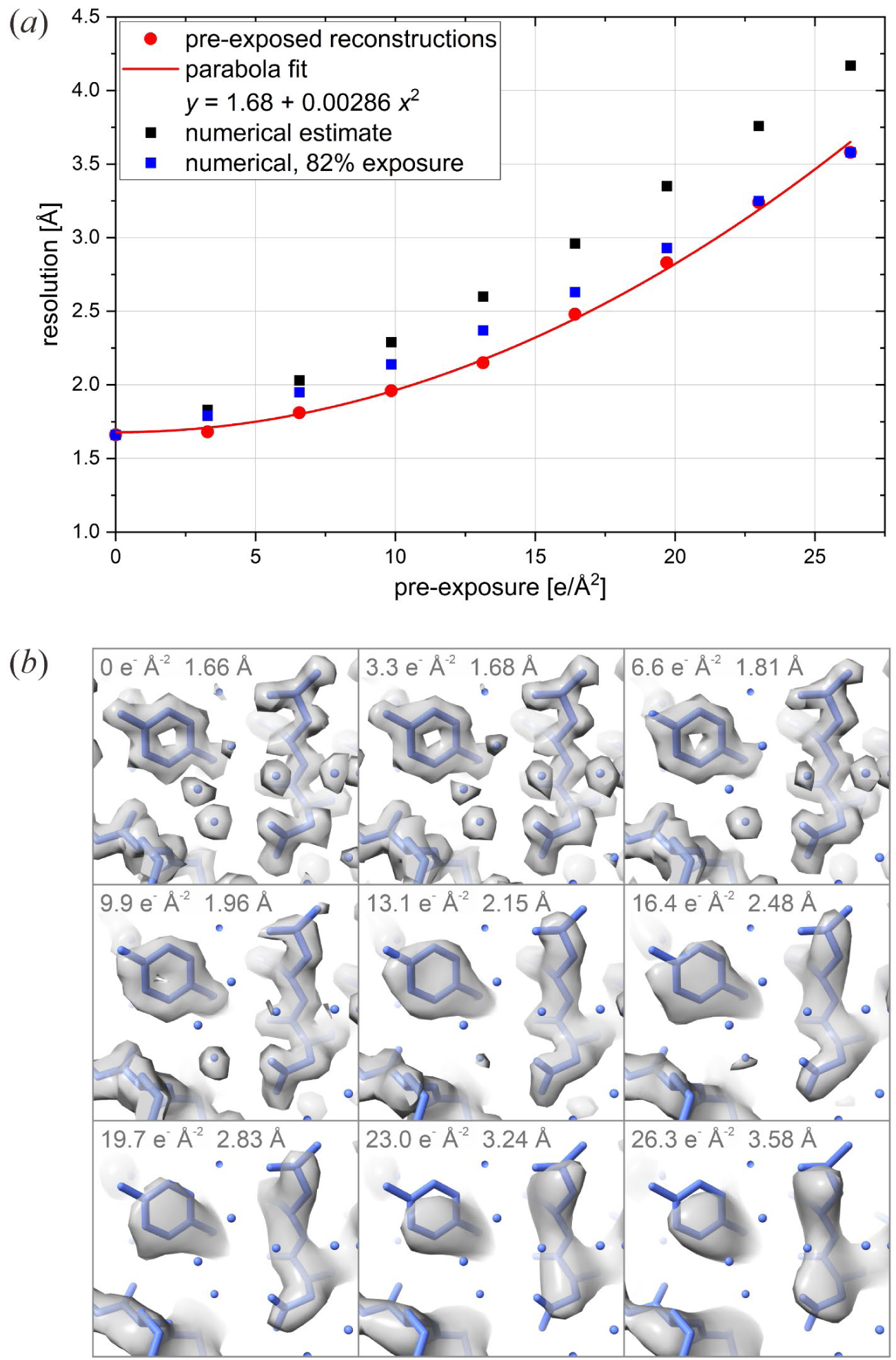
Effect of radiation damage on resolution. (*a*) Plot of the resolution of reconstructions from the first 1,000 movies containing 58k particles versus the sample pre-exposure (red symbols) by omitting a varying number of initial frames from the movies. The experimental distribution was fit well (*R*^2^ = 0.997) by a parabola (red line, coefficients in the legend). The plot also contains numerical estimates of the radiation damage effect using the measured (black symbols) and 82 % of the measured (blue symbols) experimental total exposure. (*b*) Panels illustrating map feature deterioration with increasing pre-exposure. Pre-exposure value and map resolution are shown at the top of each panel.

The reconstruction of the 100 kV dataset reached a resolution of 1.91 Å (Fig. 8). This corresponds to 123% of the physical Nyquist frequency of the detector. Map features, such as holes in aromatic sidechains, confirm the estimated resolution (Fig. 8a). Overall, this is an outstanding performance at 100 kV that closely matches what was achieved recently at 120 kV on a non-standard Glacios microscope with a narrow-gap “SP-Twin” objective lens polepiece (*C_C_* = 1.7 mm) and an Alpine (Gatan, Pleasanton, USA) camera (Chan *et al*., 2024). This configuration at 120 kV is expected to perform halfway between the Tundra microscope and the “Dublin lens” (SFEG) in Fig. 2. Having a CFEG and a narrow-gap polepiece, at 100 kV the CRYO ARM 200 II is expected to perform better than the Glacios (SFEG) at 200 kV (Fig. 2). However, the dataset did not reach higher resolution, and at this stage it is not clear what limited the practical performance. One possible reason is the relatively large pixel size and the need to rely on super-resolution processing. More advanced algorithms, such as post-acquisition super resolution (PASR) (Burton-Smith & Murata, 2023), may help to improve the result.

**Figure 8.**
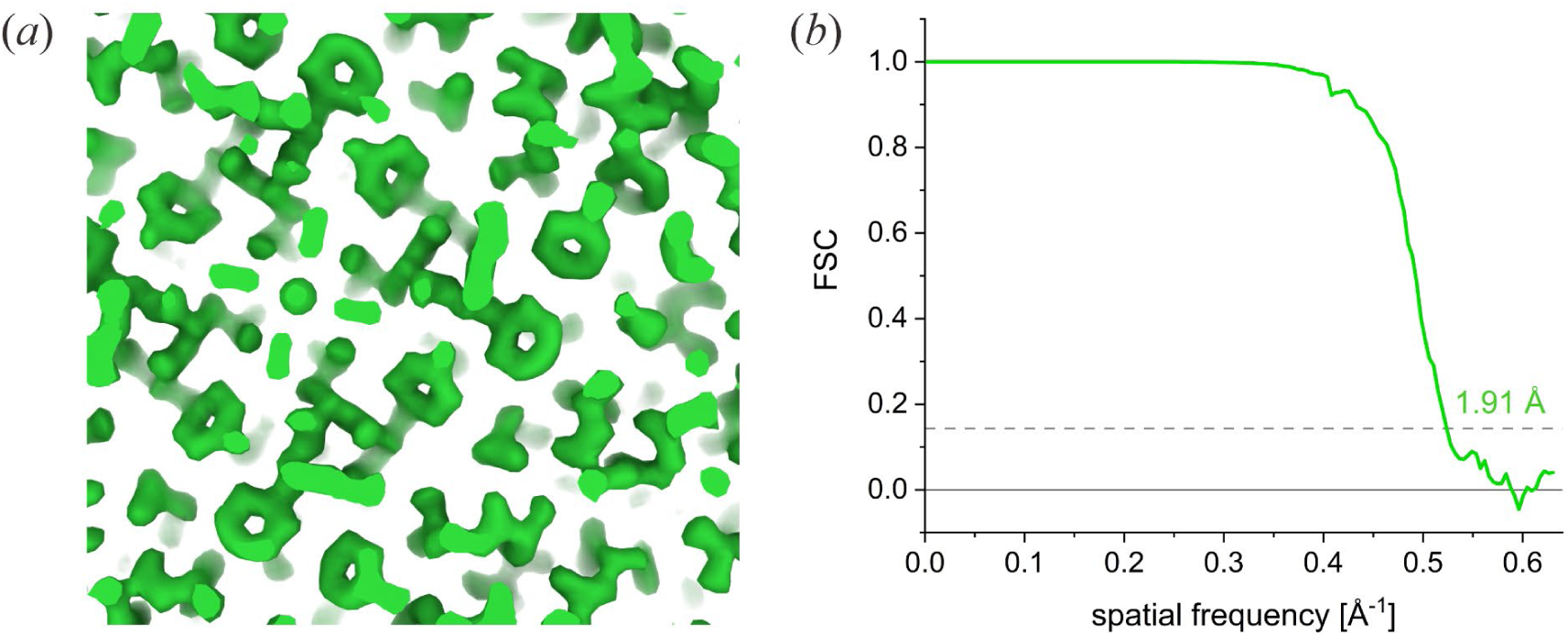
Result from the reconstruction of the 100 kV apoferritin dataset collected on the CRYO ARM 200 II with the DECTRIS SINGLA hybrid-pixel detector. The 3D reconstruction reached 1.91 Å resolution. (*a*) 3D density map showing holes in aromatic sidechains. (*b*) Gold-standard FSC plot.

## 4. Discussion

The EMDB deposition statistics (Fig. 1) indicate that despite their proven excellent performance 200 kV cryo-microscopes remain heavily underutilized. At cryo-EM facilities that also have 300 kV instruments, 200 kV microscopes are used primarily for sample screening and exploratory data collection. Understandably, having the option to use a top-of-the-line 300 kV instrument, researchers will choose to collect their data on it instead of the potentially lower-performing 200 kV machine. Such a choice was indeed justifiable with last generation 200 kV microscopes equipped with SFEG emitters which limited their performance in the sub-3 Å range, as illustrated by the lower number of 200 kV EMDB depositions in this range (Fig. 1) and the temporal envelope (Fig. 2, Glacios).

The introduction and wider adoption of CFEG emitters in recent years dramatically reshaped the cryo-EM performance landscape (Fig. 2, solid vs dashed/dotted lines). In particular, it significantly improved 200 kV performance in the 2–3 Å range (Fig. 2, Glacios vs CRYO ARM 200/Glacios 2). In addition to their optical performance advantages, CFEGs are also less costly to maintain because they do not require periodic emitter exchanges. In use, they do exhibit a gradual decline of emission current over time, necessitating tip flashes every ∼8 hours, but this is handled automatically and can be performed during cryogen filling cycles to avoid extra acquisition interruptions.

The CRYO ARM 200 II microscope tested here introduces yet another performance-enhancing feature in the form of a narrow gap objective lens polepiece, which reduces chromatic aberration and further expands the resolution envelope (Fig. 2, pink line). Its single-particle performance is comparable to top-of-the-line 300 kV instruments, both in theory and practice, as demonstrated by the results presented here (Figs. 3, 5, 6). The only practical limitation of the narrow-gap polepiece is the lack of sample tilting capability. This precludes applications such as cryo-tomography (Nogales & Mahamid, 2024), MicroED (Clabbers *et al*., 2025), or single particle samples with severe preferred orientation requiring tilted acquisition (Aiyer *et al*., 2024). Observations involving thicker samples (≥100 nm), such as cryo-tomography and 2D template matching of cellular samples, single particle analysis of viruses, liposomes, etc., will continue to benefit from higher accelerating voltages (Peet *et al*., 2019). However, for the vast majority of single-particle projects, the CRYO ARM 200 II offers uncompromising practical performance.

New technological developments, such as narrow-gap lenses and CFEGs will bring 100 kV microscopes closer to becoming the most cost-effective option for sample screening and exploratory data collection. Here, as well as in other recent works (Chan *et al*., 2024), the first sub-2 Å cryo-EM test structures from 100 kV instruments were presented. In practice, factors such as sample thickness, higher sensitivity to optical aberrations, electromagnetic disturbances, sample charging, and detector performance, must also be considered and tested before declaring success. Overall, the proof will be in the real-world sample results that will be coming out of upgraded 100 kV microscopes.

In conclusion, with the technological advances discussed here, cryo-EM instrumentation could be entering a period of experimental role hand-down, where tasks that were exclusive to 300 kV instruments will become more common on 200 kV, and those from 200 kV will be shifted to 100 kV. The goal of this process is to make cryo-EM more accessible, affordable, and prolific.

## Acknowledgements

We thank Hiromu Shinmiya (Service), Toshiyuki Yamamoto (Service), Kazuya Omoto (R&D), and Sohei Motoki (R&D) from JEOL Ltd. for their help with setting up and aligning the CRYO ARM 200 II microscope for observations at 100 kV. We are grateful to Takeyoshi Taguchi, Matthias Meffert, Saori Kawaguchi, Toshinobu Miyoshi, and colleagues from DECTRIS Ltd. for their efforts in organizing the loan and the installation of the DECTRIS SINGLA detector. We thank Yoichi Sakamaki for support in the operation and maintenance of the microscope.

## Data availability

The 100 kV and 200 kV apoferritin datasets were deposited to the Electron Microscopy Public Image Archive (EMPIAR) with accession codes EMPIAR-XXXXX and EMPIAR-XXXXXX. The 3D map and model from the 200 kV dataset were deposited to the Electron Microscopy Data Bank (EMDB) and the Protein Data Bank (PDB) with accession codes EMD-68251and PDB-22FX. The 3D map from the 100 kV dataset was deposited to the Electron Microscopy Data Bank (EMDB) with accession code EMD-68204. The full raw datasets and CryoSPARC projects are also accessible on DECTRIS CLOUD (https://www.dectris.cloud), with instant access to the workflows, including the capability to reprocess the data. For access, please contact.

