## Supplementary information for "Atomic resolution cryo-EM at 200 keV"

**Table S1.** Cryo-EM data collection, refinement, and validation

| | 200 kV dataset<br>EMPIAR-XXXXX<br>EMD-68251<br>PDB-22FX | 100 kV dataset<br>EMPIAR-XXXXX<br>EMD-68204 |
|---|---|---|
| **Data collection** | | |
| Magnification (indicated) | 150,000 | 500,000 |
| Voltage (kV) | 200 | 100 |
| Camera | Gatan K3 (CDS) | DECTRIS SINGLA |
| Energy filter slit (eV) | 20 | 20 |
| Objective aperture (μm) | - | 250 |
| Grid type | UltrAuFoil R0.6/1 | UltrAuFoil R1.2/1.3 |
| Acquisition pattern | 3 x 3 x 1 | 3 x 3 x 4 |
| Beam diameter (μm) | 0.95 | 0.53 |
| Physical pixel size (Å) | 0.3056 | 1.17 |
| Exposure rate ($e^-$ $pixel^{-1}$ $s^{-1}$) | 3.3 | 23.8 |
| Exposure time (s) | 1.51 | 3.0 |
| Electron exposure ($e^-$ Å$^{-2}$) | 53.4 | 52.2 |
| Defocus range (μm) | -0.2 – -0.8 | -0.2 – -0.8 |
| Movies (no.) | 13,654 | 17,424 |
| Frames (no.) | 81 | 13,500 |
| Fractions (no.) | 81 | 60 |
| **Data processing** | | |
| Symmetry imposed | O | O |
| Initial particle images (no.) | 652,257 | 349,779 |
| Final particle images (no.) | 615,248 | 250,077 |
| Map resolution, FSC 0.143 (Å) | 1.24 | 1.91 |
| Map resolution range (Å) | 1.22 – 1.25 | 1.85 – 1.91 |
| **Model Refinement** | | |
| Initial model used | PDB-7A4M | |
| Model composition | | |
| Non-hydrogen atoms | 1814 | |
| Protein residues | 173 | |
| Metals | 2 | |
| Water molecules | 201 | |
| R.m.s. deviations | | |
| Bond lengths (Å) | 0.0138 | |
| Bond angles (°) | 1.90 | |
| Validation | | |
| MolProbity score | 1.44 | |
| Clashscore | 5.97 | |
| Poor rotamers (%) | 1.12 | |
| CaBLAM outliers (%) | 0.6 | |
| Ramachandran plot | | |
| Favored (%) | 97.7 | |
| Allowed (%) | 2.3 | |
| Outliers (%) | 0 | |

**Table S2.** Effect of radiation damage by pre-exposure on the resolution

| Omitted initial movie frames | Corresponding pre-exposure ($e^{-}$ Å$^{-2}$) | 3D refinement resolution (Å) | Theoretical estimate (Å) | Theoretical estimate, 82% exposure (Å) |
|---|---|---|---|---|
| 0 | 0 | 1.66 | 1.66 | 1.66 |
| 5 | 3.3 | 1.68 | 1.83 | 1.79 |
| 10 | 6.6 | 1.81 | 2.03 | 1.95 |
| 15 | 9.9 | 1.96 | 2.29 | 2.14 |
| 20 | 13.1 | 2.15 | 2.60 | 2.37 |
| 25 | 16.4 | 2.48 | 2.96 | 2.63 |
| 30 | 19.7 | 2.83 | 3.35 | 2.93 |
| 35 | 23.0 | 3.24 | 3.76 | 3.25 |
| 40 | 26.3 | 3.58 | 4.17 | 3.58 |

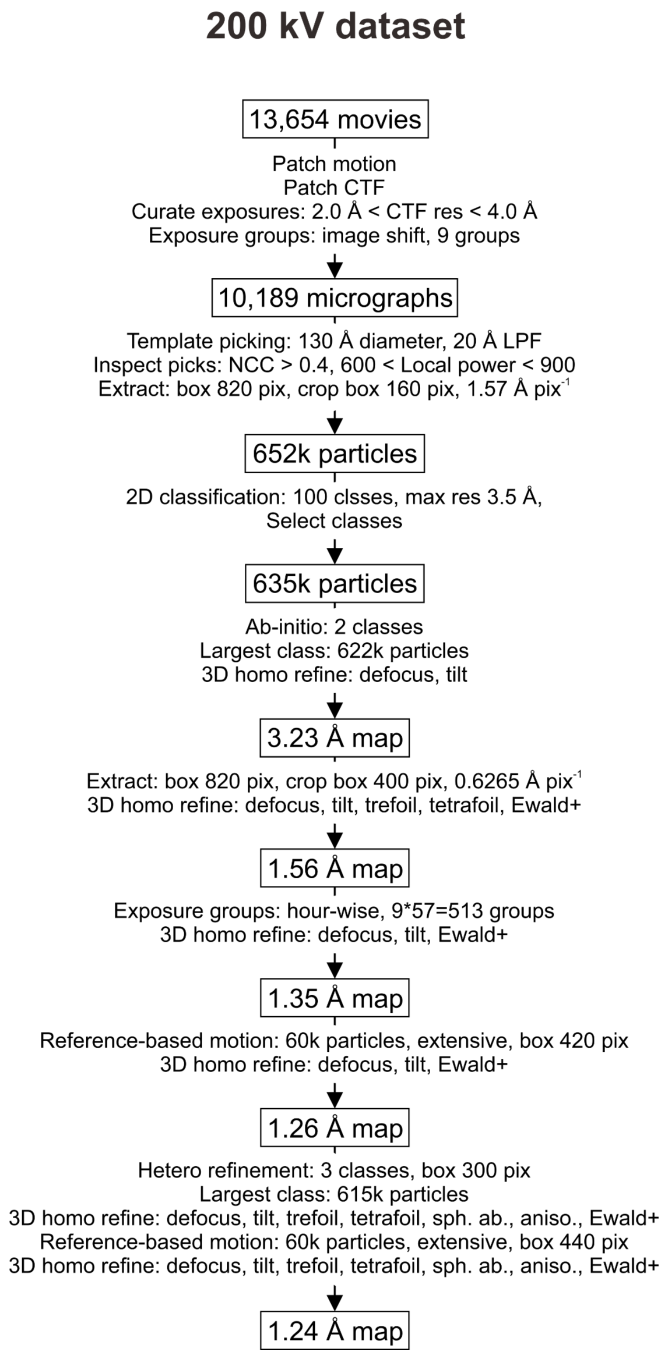


**Supplementary Figure S1.** Data processing workflow for the 200 kV dataset

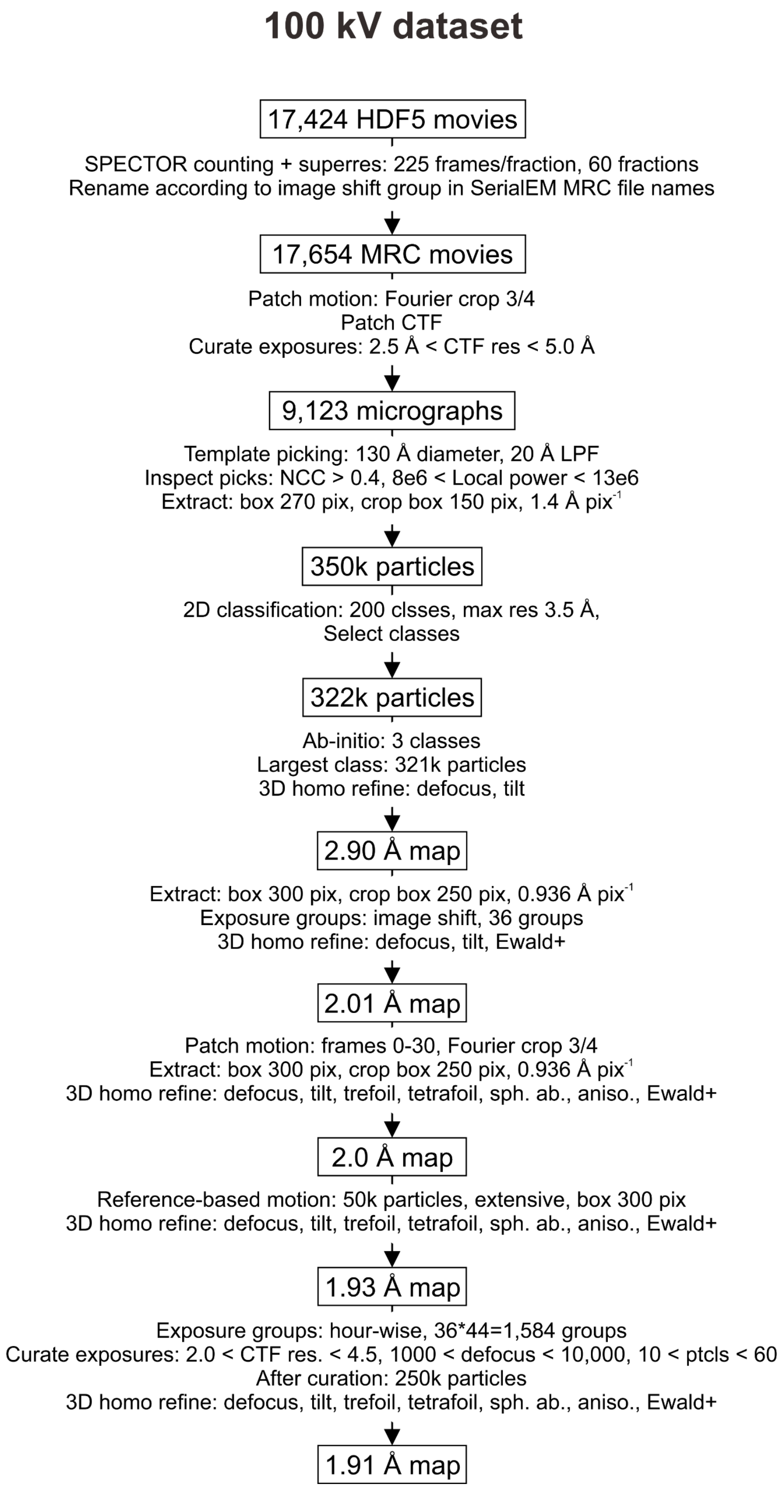


**Supplementary Figure S2.** Data processing workflow for the 100 kV dataset

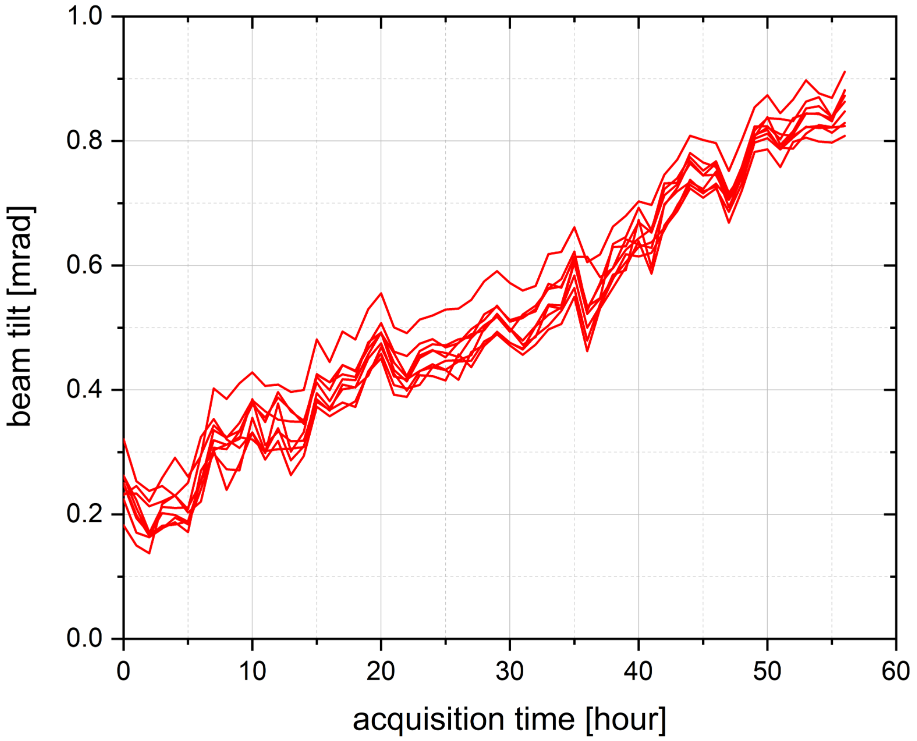


**Supplementary Figure S3.** Beam tilt evolution for the nine image shift exposure groups over the acquisition period of the 200 kV dataset.
